# A triresidue motif in the GLUTAMATE RECEPTOR-LIKE 3.3 C-tail interacts with IMPAIRED SUCROSE INDUCTION 1 and controls long distance wound signaling

**DOI:** 10.1101/2020.10.06.327924

**Authors:** Qian Wu, Stéphanie Stolz, Archana Kumari, Gaétan Glauser, Edward E. Farmer

**Affiliations:** Department of Plant Molecular Biology, Biophore, University of Lausanne, CH-1015 Lausanne, Switzerland; Plant Molecular Biology Unit, Division of Biochemical Sciences, CSIR-National Chemical Laboratory, Pune 411008, Maharashtra, India; Neuchâtel Platform of Analytical Chemistry, University of Neuchâtel, CH-2000 Neuchâtel, Switzerland

**Keywords:** wounding, phloem, jasmonate, electrical signaling

## Abstract

*Arabidopsis* Clade 3 *GLUTAMATE RECEPTOR-LIKE* (*GLRs*) genes are primary players in wound-induced electrical signaling and jasmonate-activated defense responses. As cation-permeable ion channels, previous studies have focused on resolving their gating properties and structures. However, little is known regarding to the regulatory mechanism of these channel proteins. Here, we report that the C-tail of GLR3.3 contains key elements that control its function in long distance wound signaling. GLR3.3 without its C-tail failed to rescue the *glr3.3a* mutant. To further investigate the underlying mechanism, we performed a yeast two-hybrid screen. IMPAIRED SUCROSE INDUCTION 1 (ISI1) was identified as an interactor with both the C-tail and the full-length GLR3.3 *in planta*. Reduced function *isi1* mutants had enhanced electrical activity and jasmonate-regulated defense responses. Furthermore, we found that a triresidue motif RFL (R884, F885 and L886) in the GLR3.3 C-tail is essential for interacting with ISI1. RFL mutation abolished GLR3.3 function in electrical signaling and jasmonate-mediated defense gene activation. Our study shows the importance of the C-tail in GLR3.3 function, and reveals parallels with the ipnotropic glutamate receptor regulation in animal cells.

## Introduction

Clade 3 *GLUTAMATE RECEPTOR-LIKE* (*GLR*) genes are ancient ion channels that play diverse roles in both sporophytes and gametophytes throughout the plant kingdom (1) (2). For example, these proteins function in sperm chemotaxis in a basal land plant lineage represented by the moss *Physcomitrella* (3). In the angiosperm Arabidopsis, clade 3 GLRs function in pollen tube growth (4, 5) and in lateral root development (6). One of the 7 clade 3 GLRs in *Arabidopsis*, GLR3.3, stands out for its roles in plant defense. Firstly, *glr3.3* mutations reduce the resistance of this plant to the fungus *Hyaloperonospora* (7). Secondly, along with GLR3.1, 3.2 and 3.6, GLR3.3 functions in wound-response electrical signalling leading to activation of the jasmonate pathway (8). The jasmonate pathway (9) underlies defense against many insects (10). Specifically, GLR-dependent electrical signals called slow wave potentials (SWPs) elicit the synthesis of the defense mediator jasmonoyl-isoleucine (JA-Ile) in leaves distal to wounds (8).

GLR3.3 is permeable to monovalent and divalent cations including Ca^2+^ (5), and its gating properties are currently under intense investigation (11, 12). This channel has a potentially broad gating-ligand specificity, a feature which is consistent with both *in vivo* (13) and *in vitro* studies (14). However, gating ligands may not be necessary for GLR3.3 ion channel activity when expressed in vertebrate cells (5). While it is likely that the ion channel properties of GLR3.3 are essential for its role in electrical signalling, other parts of the protein may have regulatory roles. These regions include the cytoplasmic C-tail of this protein.

Like their mammalian homologs ionotropic glutamate receptors (iGluRs), plant GLRs are predicted to be multi-membrane spanning proteins possessing a large extracellular amino terminal domain (ATD), one ligand binding domain (LBD), three transmembrane domains, one pore region and a cytoplasmic carboxyl terminal domain (CTD) (15). Despite sharing conserved structural and functional features in the LBD and pore region that are important for their channel properties, the CTDs of mammalian iGluRs and plant GLRs display variable amino acid sequences (2). GLR3.3 has an 83-amino acid C-tail which is theoretically long enough to associate physically with other proteins. Indeed, the C-tails of all clade 3 GLRs contain putative endoplasmic reticulum retention signals (2) and the C-tails of GLR3.4 and GLR3.7 bind to 14-3-3 proteins (16) (17).

We first assessed the importance of the C-tail of GLR3.3 in leaf-to-leaf electrical signal propagation, and in wound-response jasmonate pathway activation. To do this, a C-tail deletion variant of *Arabidopsis* GLR3.3 was generated. Plants harboring the *glr3.3a* reduced-function mutation (8) were transformed with this construct tagged with the fluorescent marker protein VENUS (18) and expressed under the control of the *GLR3.3* promoter. These plants were then assessed for GLR3.3 function in SWP signaling. The main pool of GLR3.3 protein detected in leaf vascular tissue is known to localise to phloem sieve element endoplasmic reticulum (19). Using a recently developed primary vein extraction technique (20), the cellular localization of a GLR3.3 C-tail truncation variant was examined. In parallel, we conducted a yeast two hybrid screen against the GLR3.3 C-tail. And we identified a putative interactor with the C-tail and further characterized its function in long distance wound signaling.

## Results

### The C-tail of GLR3.3 is required for its function

Despite of functional redundancy among *GLR3.3, GLR3.6 and GLR3.1*, the sequence alignments showed their C-tail sequences are variable with the exception of several conserved residues (Fig. 1*A*). To determine if the entire C-tail of GLR3.3 was essential for function, we generated GLR3.3 lacking its C-tail. The truncated GLR3.3 gene was then fused with VENUS and transformed into the *glr3.3a* mutant background in order to investigate its ability to rescue the *glr3.3a* mutant phenotype. SWPs in leaves distal to wounds were measured in two independent lines of the C-tail deleted plants, along with wild type (Col-0), a *glr3.3a* mutant, and the full-length (FL) complemented line (*GLR3.3pro:GLR3.3-VENUS#2.3*) (19) as controls. No substantial differences were found in the amplitude of the SWPs recorded from all the genotypes (Fig. 1*B*, left panel). However, compared to the wild type and the full-length complemented plants, the lines which express GLR3.3 lacking its C-tail (*ΔCT-10-6b#* and *ΔCT-18-5a#*) failed to rescue the *glr3.3a* knockout phenotype and showed strongly reduced, *glr3.3a* mutant-like duration of SWPs (Fig. 1*B*, right panel). Then we tested the wound-induced *JAZ10* defense marker expression levels in these plants. *JAZ10* expression was similarly attenuated in the C-tail variants as in the *glr3.3a* mutants (Fig. 1*C*). Together, these results showed that the short C-tail of GLR3.3 is required for its functions in propagating surface potentials and in distal defense-related gene expression.

**Fig. 1.**
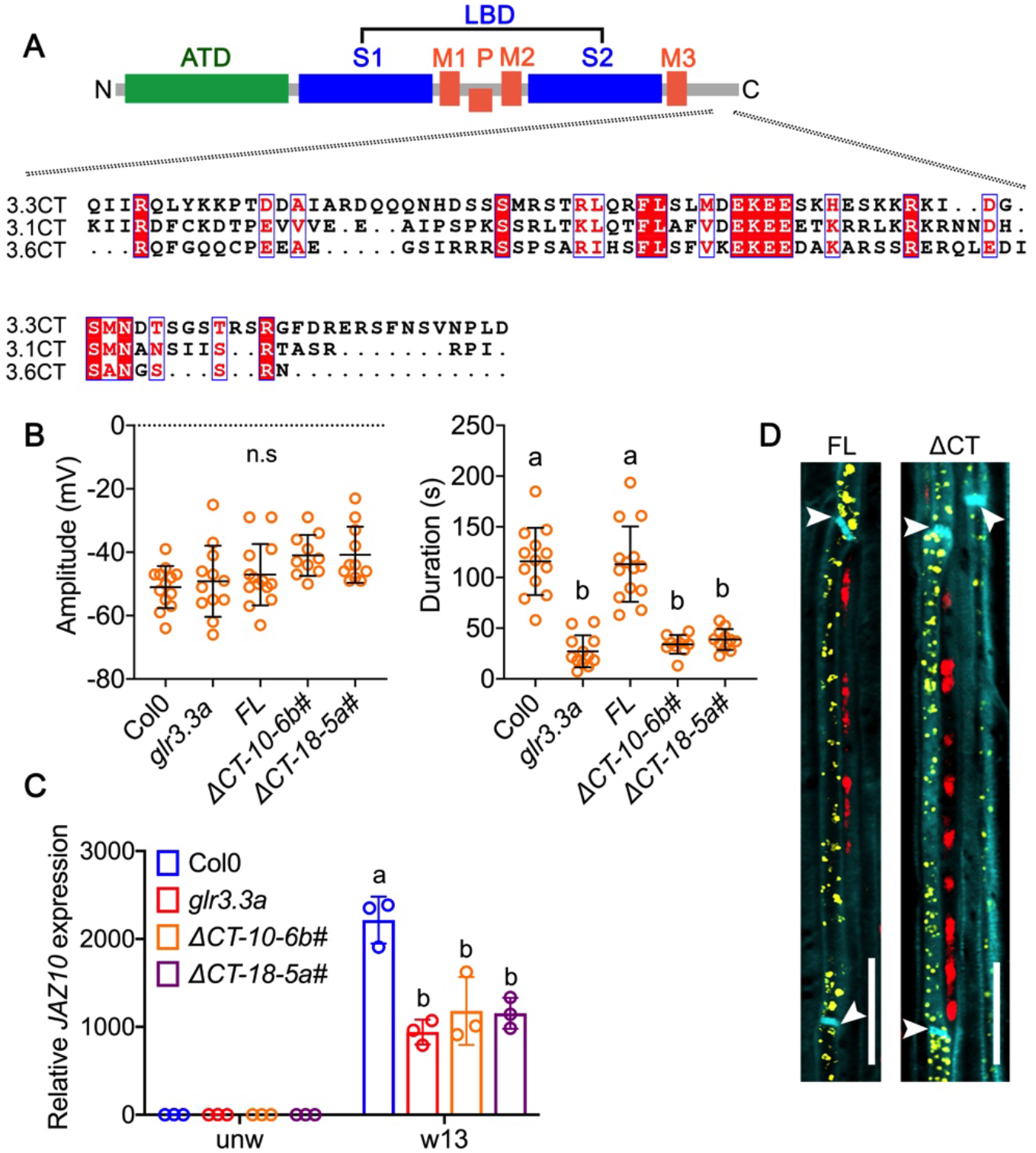
Deletion of the C-tail of GLR3.3 impairs its function in wound-induced electrical signaling and defense activation, but not subcellular distribution. (*A*) Schematic diagram showing GLR architecture and sequence alignments of the C-terminal tails from three clade 3 GLRs. ATD: amino-terminal domain. LBD: ligand binding domain. S1 and S2: segment 1 and segment 2. M1 to M3: membrane-spanning domain 1 to 3. P: pore region. CT: C-tail. (*B*) Amplitude and duration of surface potential changes on leaf 13 after wounding leaf 8. Wild type, *glr3.3a* mutants, *glr3.3* complemented plants, as well as two independent lines for the C-tail deletion plants were measured. The orange points represent individual plants. n=10-14. The horizontal bars indicate the mean values. Error bars show S.D. The different letters indicate significant differences (*P*<0.01) after one-way ANOVA. n.s, not significant. FL: Full length. ΔCT is the deletion form of the GLR3.3 C-tail from residues 850 to 933. (*C*) *JAZ10* expression levels in unwounded leaf 8 and distal leaf 13 after wounding leaf 8. Data shown are means ± SD. Each colored circle represents one biological replicate. n=3. The different letters indicate significant differences (*P*<0.01) after two-way ANOVA. (*D*) Subcellular localization of full-length GLR3.3 protein and its C-tail deletion. Midveins from plants expressing GLR3.3-VENUS (FL) or GLR3.3 ΔCT–VENUS (ΔCT) fusions under the *GLR3.3* promoter were extracted and VENUS was localized by confocal microscopy. Yellow is signal from GLR3.3-VENUS. Red is chlorophyll autofluorescence from companion cells. Cyan marks outlines of the cells. Arrowheads indicate the positions of sieve plates. Images were taken with the same parameters. Bars=20 μm.

We next investigated whether deletion of the GLR3.3 C-tail altered its cellular distribution, thus leading to loss function of the protein. To test this, we compared the expression of the truncated GLR3.3-VENUS fusion protein with the full-length GLR3.3 fusion. As reported by Nguyen et al. (2018) and also found in this work, the major pool of GLR3.3-VENUS were localized to ER-like structures in sieve elements (Fig. 1*D*, FL). Interestingly, a similar VENUS distribution pattern was obtained in plants expressing the GLR3.3 variant lacking its C-tail (Fig. 1*D*, ΔCT). Given that neither the expression nor the localization of GLR3.3 was impaired, the inability of GLR3.3 C-tail deletion to rescue the *glr3.3a* phenotype indicates that the C-tail of GLR3.3 contains motifs that are functionally important to preserve GLR protein activity or to interact with other regulators.

### ISI1 interacts directly with GLR3.3 *in vivo*

To further explore the role of the C-tail of GLR3.3 in wound-induced electrical signaling, a yeast two-hybrid (Y2H) screen was conducted using the C-tail as bait against a cDNA library from Arabidopsis rosette leaves. This screen aimed at finding interacting proteins that could potentially regulate GLR3.3 function. The full list of all the candidate genes from the screen can be found in the Supplementary Dataset 1. *IMPAIRED SUCROSE INDUCTION 1* (*ISI1*) was the candidate with the highest confidence for interactions. *ISI1* encodes a highly conserved plant-specific protein with structural similarities to Arm repeat proteins (21). Loss-of-function *isi1* mutants were reported to display impaired sucrose induction of starch biosynthetic genes (21). Beyond this, little is known regarding the biological function of ISI1. However the *ISI1* promoter was reported to be active in the phloem of adult-phase leaves (21). *GLR3.3* is also expressed in the phloem (19). The interaction between ISI1 and the C-tail of GLR3.3 (3.3CT) was retested in the yeast system. Indeed, ISI1 was able to interact with GLR3.3CT, but not with either the C-tail of GLR3.1 (3.1CT) or GLR3.6 (3.6CT) (Fig. 2*A*). To further confirm their interaction *in vivo*, we took advantage of the firefly luciferase complementation imaging (LCI) system to transiently express different combinations of cLUC and nLUC fusion proteins in *N. benthamiana* leaves. In contrast to the negative controls, ISI1 interacted with both the C-tail and the full-length GLR3.3 proteins (Fig. 2*B*). ISI1 is therefore a new interactor of GLR3.3 *in planta*.

**Fig. 2.**
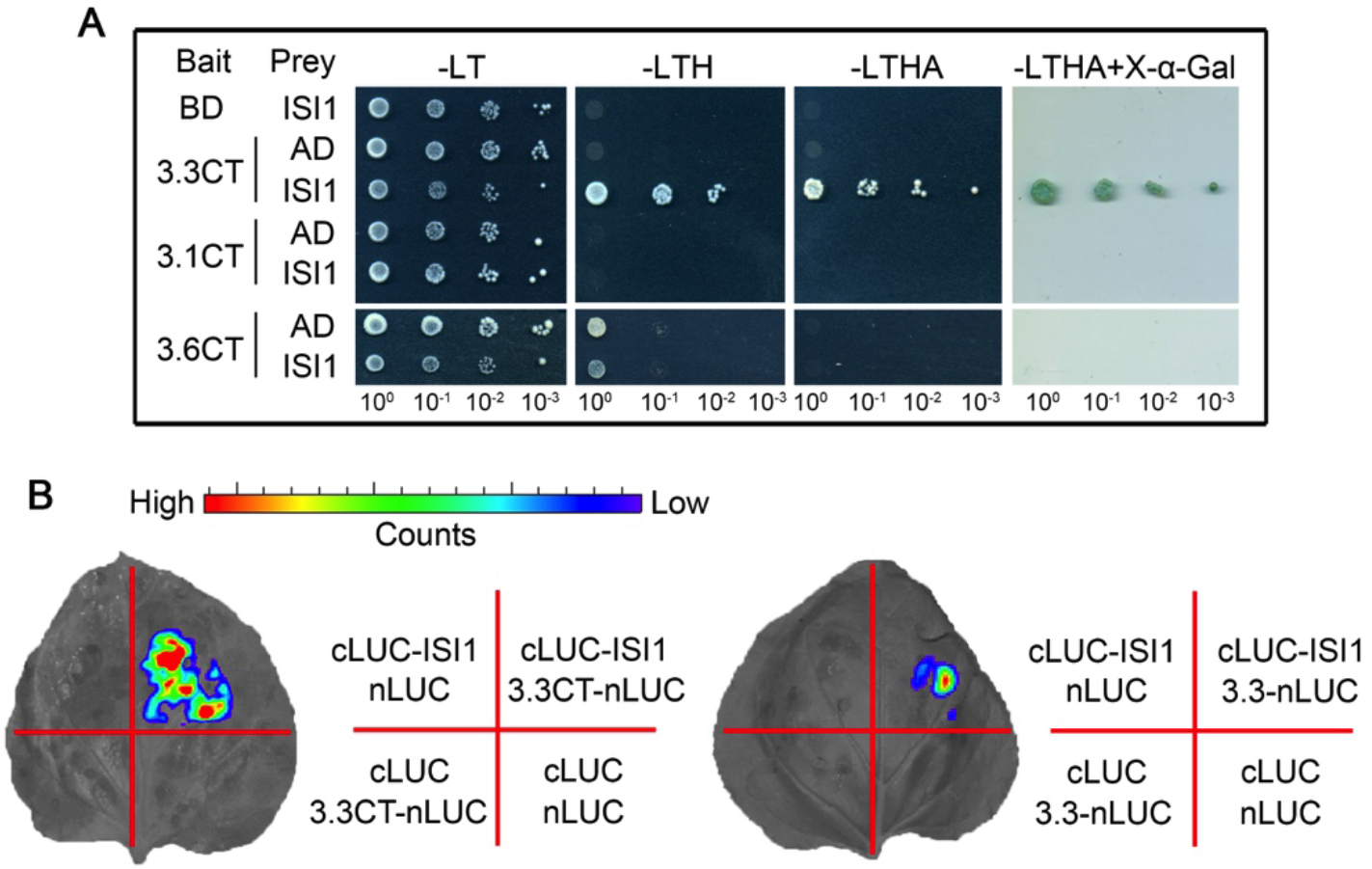
ISI1 interacts with the GLR3.3 C-tail *in vivo*. (*A*) ISI1 interacts specifically with the C-tail of GLR3.3, but not GLR3.1 and GLR3.6 in yeast two hybrid (Y2H) assays. AD-fused ISI1 was co-transformed with BD-fused C-tail of GLR3.3, GLR3.1 or GLR3.6. Empty AD or BD vectors were included as negative controls. The interactions were tested by growing yeast cells on different selective media. Photographs were taken after 3 days for yeast grown on Leu-Trp- (-LT)/Yeast Nitrogen Base (YNB) medium or 5 days for the other yeast groups on selective Leu-Trp-His- (-LTH), Leu-Trp-His-Ade (-LTHA) and –LTHA plus X-α-gal media. (*B*) Firefly Luciferase complementation imaging (LCI) assay showing the interactions of ISI1 with the C-tail and the full-length GLR3.3 proteins. Construct pairs as indicated in the right panel of (*B*) were coexpressed in *N. benthamiana* leaves. Representative pictures are shown for the interactions. Color scale reflects LUC activity.

### Vasculature-associated *ISI1* is involved in wound-Induced long distance signaling

The physical interaction between GLR3.3 and ISI1 implied that ISI1 might be involved in long distance wound responses. Firstly we determined if ISI1 protein was produced in the tissues where GLR3.3 was found. We created translational fusion plants expressing *IS11_pro_:ISI1-GUSPlus* in the WT background. In 3-week-old rosettes, ISI1-GUS was distributed ubiquitously in the whole leaf, including the trichomes (Fig. 3*A* and *B*). Then in order to investigate the cell types that ISI1-GUS is associated with, a transversal section was generated from the petiole of an expanded leaf. GUS staining was detected in cells in both the phloem and xylem regions (Fig. 3*C*), and ISI1-GUS appeared to be more abundant in the vascular bundles than in the surrounding cells (Fig. 3*C*). To investigate if ISI1 played a role in wound signaling, two T-DNA insertion alleles of the *ISI1* gene, *isi1-2* (Salk_014132) (21) and *isi1-3* (Salk_045849) were obtained (Supplementary Fig. 1*A* and *B*). Wound-induced distal SWPs were measured in these two lines. Compared to WT, prolonged durations of the surface potential were detected in these two *isi1* alleles without affecting the signal amplitudes (Fig. 3*D* and *E*, Supplementary Fig. 1 *C* and *D*). The *isi1-2* allele was chosen for detailed analyses. This mutant was crossed with *glr3.3a* in order to study their genetic interactions. Without visible differences in amplitude, the *glr3.3a* mutation suppressed the effect of *isi1* on wound-activated SWPs in terms of duration in the *isi1-2 glr3.3a* double mutants (Fig. 3*D* and *E*), indicating that ISI1 functions through a GLR3.3-dependent pathway.

**Fig. 3.**
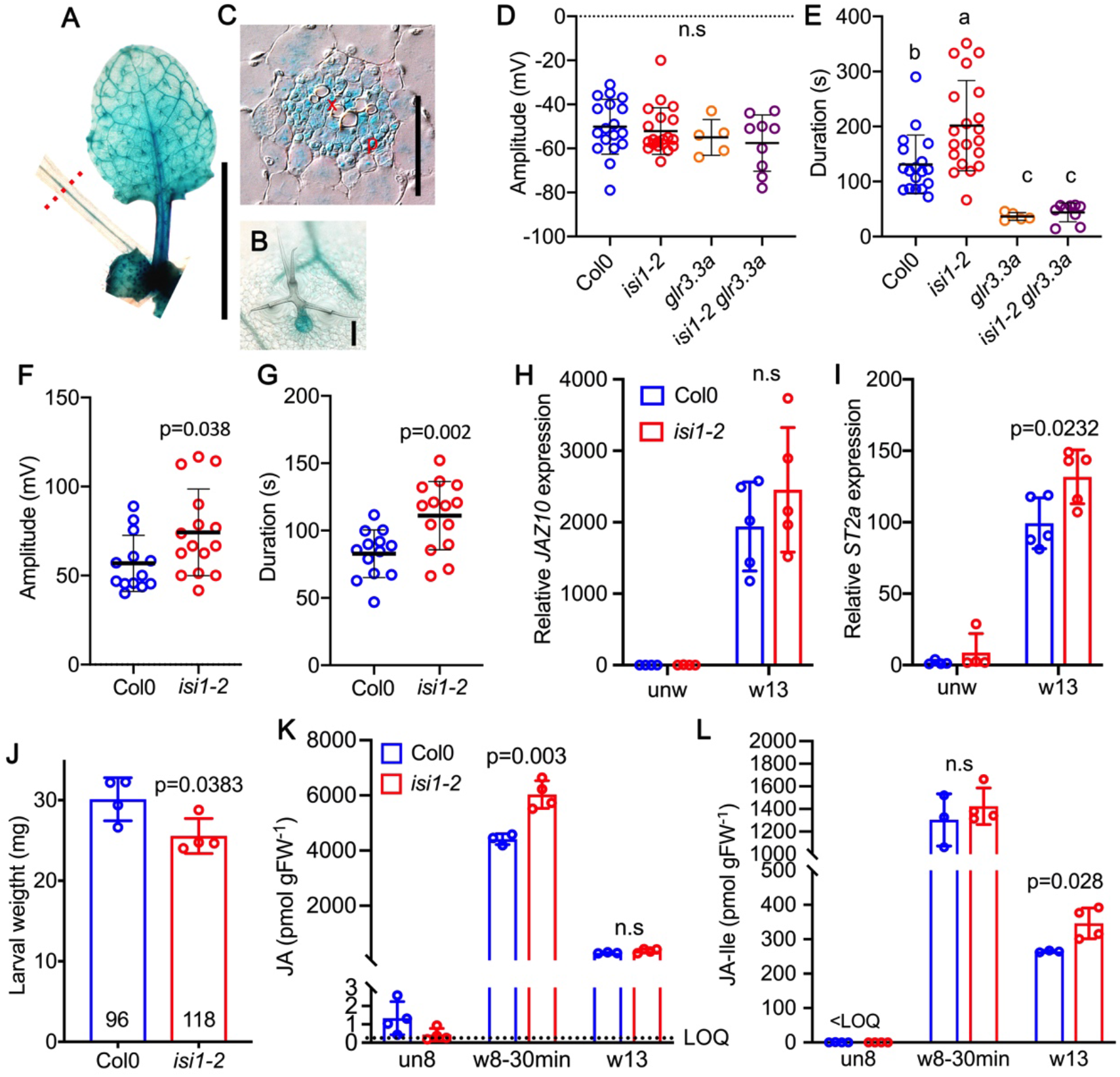
Vasculature-associated *ISI1* affects surface potentials and sieve elements-specific electrical signal durations, as well as defense. (*A*-*C*) GUS staining for ISI protein expression pattern from three-week-old *ISI1_pro_:ISI1_genomic_-GUS* plants. GUS activity for a rosette leaf is shown in (*A*). Bar=1 cm. (*B*) Stained trichome, bar=100 μm. (*C*) Transversal section of the petiole at the position indicated by the red dashed line in (*A*). Bar=50 μm. P, phloem region. X, xylem region. (*D* and *E*) Amplitude (*D*) and duration (*E*) of surface potential changes on leaf 13 after wounding leaf 8 recorded from different genotypes. Colored circles indicate individual biological replicates. n=5-20. The horizontal bars indicate the mean values. Error bars show S.D. The different letters indicate significant differences after one-way ANOVA. n.s, not significant. (*F* and *G*) Amplitudes (*F*) and durations (*G*) for sieve elements-specific electrical signals detected with aphid electrodes from leaf 13 after wounding leaf 8. Colored circles shown are measurements from individual plants. n=13-14. The horizontal bars indicate the mean values. Error bars show S.D. *p* values were calculated with two-tailed Student’s *t*-tests. (*H* and *I*) Expressions of the jasmonate response genes *JAZ10 (H)* and *ST2a* (*I*) in distal leaf 13 after wounding leaf 8. n=4-5. (*J*) Larval weight gain on *isi1-2* versus wild type plants after feeding for 11 days. Four biological replicates were analyzed. Each replicate is the average larval weight from 44 larvae that were initially placed in one box. Numbers in each bar indicate how many larvae survived in the end of the experiment. (*K* and *L*) JA (*K*) and JA-Ile (*L*) contents in *isi1-2* in comparison with the WT plants in both wounded and distal leaves. LOQ: limit of quantification. Data shown are means ± SD. n=3-4. *p*-values were calculated with two-tailed Student’s *t*-tests. n.s, not significant.

To further test if mutation in *ISI1* was responsible for the altered wound-induced electrical activity, the *isi1-2* mutant was complemented with an *ISI1*-encoding genomic fragment to which a fluorescent mCherry tag was added to its C-terminus. The prolonged duration of electrical signals in *isi1-2* upon wounding was restored to wild type level by driving *ISI1* expression in this line (Supplementary Fig. 1*E* and *F*). Moreover, consistent with the ISI1-GUS expression pattern, western blot analysis with this complemented line showed more ISI1-mCherry fusion protein was detected in midvein protein extracts than in the extracts from leaf lamina without midvein (Supplementary Fig. 1*G*), reinforcing the likely roles of ISI1 in vascular tissues. Then we examined phloem electrical signals using an electrical penetration graph (EPG) approach that employs living aphids as electrodes to measure electrical activity in sieve elements (22). Leaf 8 was wounded when the aphids were in the phloem-feeding phase in the sieve elements (SEs) of leaf 13. The resulting electrical signals in SEs in leaf 13 were recorded from both wild type and *isi1-2* plants. Then we quantified the amplitudes as well as the overall durations of the depolarization phases. As shown in Fig. 3*F* and *G*, compared to WT, *isi1-2* increased both the amplitudes and durations of the depolarization signal. The prolonged duration of sieve element electrical signals in *isi1-2* is consistent with that measured with surface electrodes, confirming the roles of *ISI1* in wound-induced electrical events.

Previously a strong correlation between the duration of SWPs and the activation of jasmonate-mediated defense was observed in *glr* mutants, where a shorter duration corresponded with reduction of defense marker gene expression (8, 19). Also, in loss-of-function *aha1* proton pump alleles, a prolonged duration of SWPs is coupled with increased defense responses (23). In our experiments, the expression of the defense marker gene *JAZ10* was weakly but not significantly more induced in *isi1-2* relative to the WT (Fig. 3*H*). We then performed mRNA sequencing with leaves distal to wounds in *isi1-2* and WT plants, in order to assess the impact of ISI1 on the wound response. 1452 genes, which represented 48% of all the increased transcripts (adjusted *p*<0.05), were upregulated more than twofold in *isi1-2* as well as WT plants (Supplementary Fig. 2). Among all the differentially expressed genes (adjusted *p*<0.05), those that were more induced in *isi1-2* were further classified by Gene Ontology analysis using GO term biological processes. “response to jasmonate” and “response to wounding” were among the most significantly overrepresented genes expressed to higher levels in *isi1-2* than WT (Supplementary Fig. 3), suggesting ISI1 primarily affects the branch of jasmonate-regulated genes in response to wound. One of the genes of interest was *ST2a*, which encodes a sulfotransferase and is responsive to jasmonate (24). We retested this gene and found that its transcripts were more accumulated in wounded *isi1-2* than wounded WT (Fig. 3*I*). In addition, insect resistance was assessed on *isi1-2* and WT plants. After feeding for 11 days, *Spodoptera littoralis* gained less weight on *isi1-2* relative to WT (Fig. 3*J*). In agreement, jasmonic acid (JA) and the precursor 12-oxo-phytodienoic acid (OPDA) were accumulated to higher levels in the wounded leaves of *isi1-2* than in the WT (Fig. 3*K* and Supplementary Fig. 4). An increased amount of jasmonoyl-isoleucine (JA-Ile) was also found in distal leaves of *isi1-2* compared to WT (Fig. 3*L*). Collectively, our findings support roles for ISI1 in various wound-associated responses.

### ISI1 binding sites in the GLR3.3 C-tail are required for GLR3.3 function

Next, in an attempt to identify the binding sites of ISI1 and GLR3.3CT, three serially truncated versions of GLR3.3 C-tail (3.3CT) were constructed as baits: 3.3CT (850-913 amino acids (aa)), 3.3CT (850-903 aa) and 3.3CT (850-883 aa) (Fig. 4*A*). Each bait was co-transformed individually with the prey ISI1 protein and tested in Y2H assays. In contrast to 3.3CT (850-913 aa) and 3.3CT (850-903 aa), which were both able to bind ISI1, the shorter 3.3CT truncation (850-883 aa) lost its interaction with ISI1 (Supplementary Fig. 5*A*), suggesting the interacting residues are probably located in region 883-903 aa. To further narrow down the sites, we performed site-directed mutagenesis in this area and created different baits carrying 3.3CT point mutations. Intriguingly, among all the mutated C-tail variants (Fig. 4*B* and Supplementary Fig. 5*A*), only those carrying mutations in three amino acids Arg884 (R884), Phe885 (F885) and Leu886 (L886) (mRFL, mR and mFL) impaired the interaction with ISI1. A mutation in Ser887 (mS) next to RFL, or in a combination of 4 amino acids Lys900 (K900)/Lys901 (K901)/Arg902 (R902)/Lys903 (K903) (KKRK) did not affect their interactions (Fig. 4*B* and Supplementary Fig. 5*A*). A similar mapping strategy was applied to ISI1 in order to find the GLR3.3 C-tail interacting region. However, none of the ISI1 truncations were capable of interacting with the GLR3.3 C-tail in yeast cells, suggesting that full-length ISI1 is required for its binding to GLR3.3 (Supplementary Fig. 5*B* and *C*).

**Fig. 4.**
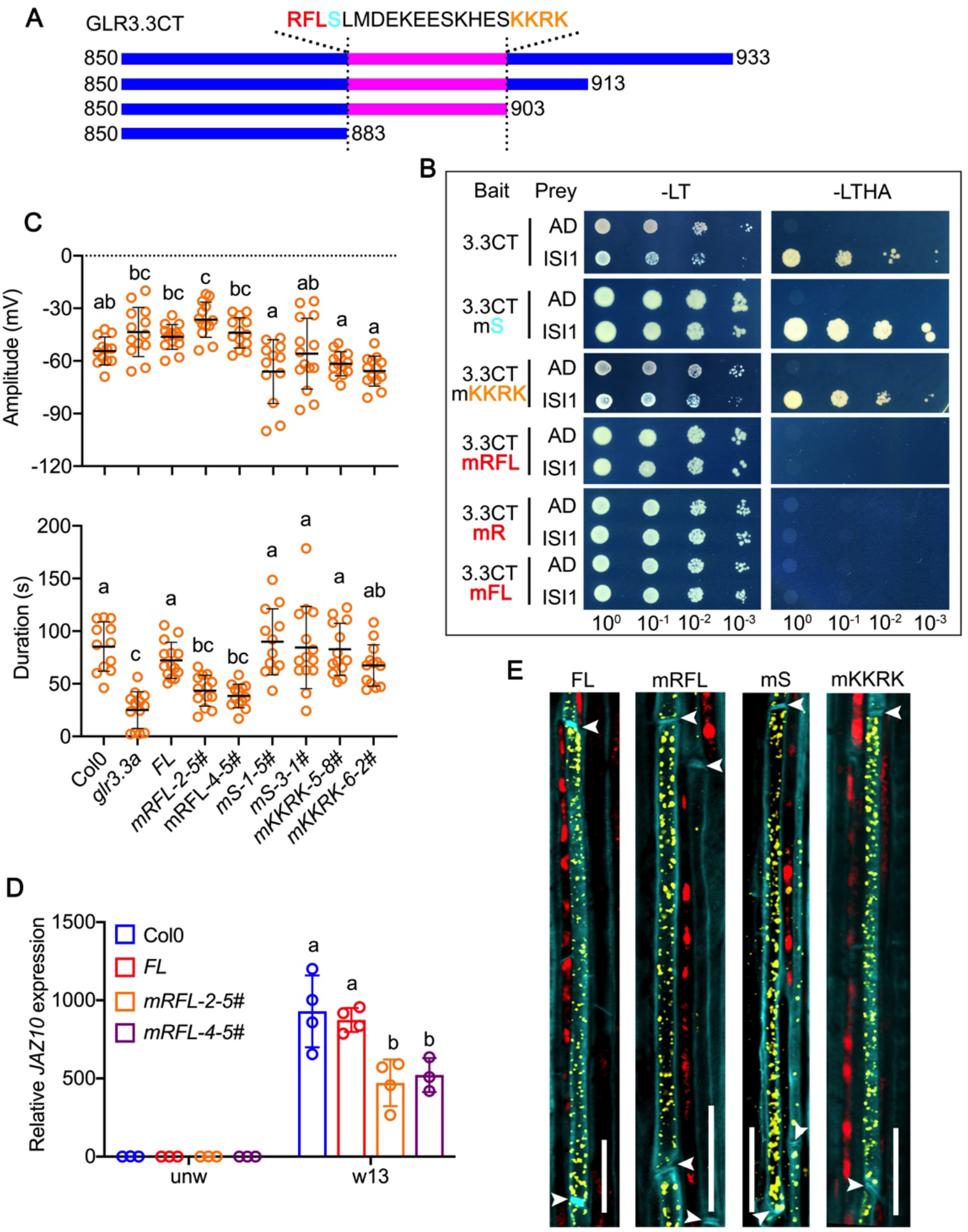
RFL residues in the GLR3.3 C-tail are required for its interaction with ISI1 and for GLR3.3 function. (*A*) Schematic diagram showing serial truncations of GLR3.3 C-tail that were used for mapping interacting sites. Amino acid sequences from position 884 to position 903 are presented. (*B*) Mutations in RFL residues (Red in (*A*)), but not S (Cyan in (*A*)) or KKRK (Orange in (*A*)), abolish its interaction with ISI1 in yeast. GLR3.3 C-tail carrying mutations in residues R, FL, RFL, S or KKRK as indicated by different colors in (*A*) were cotransformed with ISI1 into yeast cells to test the interactions. (*C*) SWPs (Amplitude and Duration) recorded in distal leaves after wounding one leaf. The yellow circles represent individual measurements. n=9-15. Two independent lines for different GLR3.3 C-tail variants were measured in comparison with WT, *glr3.3a* mutants and *glr3.3a* complemented plants. The horizontal bars indicate the mean values. Error bars show S.D. Letters represent significant differences (*P*<0.05) after one-way ANOVA. (*D*) *JAZ10* expression in distal leaves 13 in comparison with unwounded leaves. mRFL variant was analyzed compared to control plants. Data shown are means ± SD. Colored points indicate different biological replicates. n=3-4. The different letters indicate significant differences (*P*<0.01) after two-way ANOVA. (*E*) Subcellular distribution of GLR3.3 C-tail variants in comparison with the full-length version (FL). Yellow represents VENUS signals. Red indicates chlorophyll autofluorescence. Cyan marks the outlines of the cells. Arrowheads show the positions of the sieve plates. Bars=20 μm.

Given that RFL residues in the GLR3.3 C-tail are essential for its interaction with ISI1, we next sought to determine if these three amino acids were important for GLR3.3 biological function. To address this question, we generated plants that expressed GLR3.3 with mutations in RFL (*mRFL-2-5#* and *mRFL-4-5#*). Meanwhile, two other GLR3.3 C-tail variants were produced as controls. One had a point mutation in the Ser887 residue (*mS-1-5#* and *mS-3-1#*). The other one carried mutations in the 4 residues KKRK (3: *mKKRK-5-8#* and *mKKRK-6-2#*). Neither Ser887 not KKRK is involved in the interaction with ISI. The above GLR3.3 C-tail derivatives were fused with a VENUS tag and transformed into *glr3.3a* mutant backgrounds. Surface potentials were measured in all the plant lines. The amplitudes of the signal varied slightly among all the lines measured (Fig. 4*C*, *upper panel*). However, compared to the WT and the full-length (FL) complemented plants, plants carrying RFL mutations were incapable of rescuing the short-duration SWP detected in the *glr3.3a* mutant. In contrast, the Ser887 point mutation plants and the mKKRK plants all complemented the electrical signal phenotype of *glr3.3a* (Fig. 4*C*, *lower panel*). Next, distal *JAZ10* expression in mRFL plants which have compromised electrical signals was studied. Consistently, *JAZ10* induction in distal leaf 13 upon wounding leaf 8 was attenuated in mRFL plants in comparison to WT and the full-length complemented line (Fig. 4*D*). Finally, despite the different capacities in rescuing the *glr3.3a* electrical signaling phenotypes, all the GLR3.3 variants above showed similar subcellular distribution in sieve elements as the full-length protein (Fig. 4*E*).

## Discussion

Vertebrate ionotropic glutamate receptors (iGluRs) and plant GLRs are related. The C-tails of iGluRs can play critical roles in receptor localisation and function and can bind a variety of regulatory proteins that act to optimize their function. The importance of cytoplasmic C-termini of vertebrate relatives of the plant GLRs is exemplified in synaptic iGluRs. Vertebrate iGluRs function not only as ligand-gated ion channels, but they can also form large signalling complexes with diverse proteins. For example, vertebrate NMDA receptors in postsynaptic membranes are organized in signalling modules of over 1 MDa (25). In these complexes, the cytoplasmic C-tails of the NMDA receptors bind directly and indirectly to multiple partners including scaffold proteins, protein kinases, protein phosphatases, and transcriptional corepressors (26–28). Furthermore, the C-tail of vertebrate GluN1 can itself translocate to the nucleus to regulate synapse function (29). While many vertebrate NMDA receptor subunits such as AMPA receptors have long (>500 amino acids) cytoplasmic C-tails, other iGluRs from vertebrates with smaller C-tails also form complexes with unrelated proteins (30). Proteins with a potential to interact with the GLR3.3 C-tail have not, to our knowledge, been identified. To initiate this study we first tested the effect of GLR3.3 C-tail truncation *in planta*. The major pools of GLR3.3 in the vasculature localise to the endoplasmic reticulum in sieve elements (19). GLR3.3 lacking its entire C-tail did not complement the *glr3.3a* reduced function mutant (Fig. 1*B-C*) and we assumed that this might have been due to mislocalisation of the protein. However, the cellular and subcellular localization of the C-tail deletion variant was similar to that of the WT protein (Fig. 1*D*). Given the essential role of the GLR3.3 C-tail *in planta* and the fact that the C-tails of other GLRs can bind 14-3-3 proteins (17, 31), we initiated a Y2H screen using the entire GLR3.3 C-tail as a bait. The primary candidate obtained was the IMPAIRED SUCROSE-INDUCTION (ISI1; At4g27750), a plant-specific protein that was reported to act in the phloem as a positive regulator of the expression of sucrose-inducible genes (21).

We confirmed the GLR3.3 C-tail/ISI1 interaction *in planta* (Fig. 2). Their physical interaction promoted us to investigate whether ISI1 is involved in wound responses. Both phloem and xylem tissues have been shown to be crucial to transmit SWPs(19) (32). The Rook et al. (2006) study reported ISI1 promoter activity in the phloem. A translational reporter produced in our study showed ISI1 protein in xylem- and phloem-associated cells as well as in trichomes (Fig. 3*A-C*). We noted that the expression pattern of ISI1 was similar to that observed for the *Arabidopsis H^+^-ATPase 1* (*AHA1*) gene (23). AHA1 functions in the SWP repolarisation phase and in the regulation of jasmonate synthesis in response to wounding (23). We first tested whether *isi1-2* mutants affected leaf-to-leaf electrical signalling. Similar to the reduced function *aha1-7* mutant (23), the duration of the SWP was increased in *isi1* loss-of-function mutants, whereas signal amplitude was not affected (Fig. 3*D* and *E*, Supplementary Fig. 1 *C* and *D*). Similar results were obtained for *isi1-2* when sieve element-specific electrical signals were measured using aphid electrodes (Fig. 3*F* and *G*). The elevated electrical activity in *isi1-2* was attenuated by *glr3.3* mutation in the *isi1-2 glr3.3a* double mutant background, suggesting that these two genes interact genetically (Fig. 3*E*). Although the expression level of *JAZ10* was not significantly higher in *isi1-2* relative to WT (Fig. 3*H*), our RNAseq analysis with wounded *isi1-2* mutants showed that ISI1 affects the expression of other jasmonate inducible genes, including *ST2a*, as was confirmed in an independent experiment (Fig. 3*I*, Supplementary Fig. 2 and 3). From previous work, loss-of-function mutants in *ISI1* reduced the sucrose-induced expression of the ADP-glucose pyrophosphorylase subunit *APL3* (At4g39210; (21, 33)). We did not detect deregulation of this gene in our experiments. Additionally, *isi1-2* was more resistant to insect feeding and accumulated higher jasmonate levels than the WT after wounding (Fig. 3*J-L*, Supplementary Fig. 4), indicating that ISI1 plays a role in several aspects of wound responses. However, how ISI1 functions in jasmonate pathway induction after wounding is unclear; there is at present no indication that ISI1 is part of the canonical jasmonate pathway. It will be interesting to further investigate how the physical interaction between ISI1 and GLR3.3 accounts for ISI1’s function in wound signaling.

Next we mapped the binding site of ISI1 on the GLR3.3 C-tail using Y2H assays. This revealed that ISI1 bound to a trio of residues (RFL) in the central region of the C-tail (Fig. 4*B* and Supplementary Fig. 5). Mutation of these residues abolished GLR3.3 function. In contrast, GLR3.3 with Ser887 mutation or KKRK mutations located from 900-903 aa, which are still capable in interacting with ISI1 in yeast, fully complements the *glr3.3a* mutant phenotypes (Fig. 4*C-D*). This led us to speculate that the binding of ISI1 to GLR3.3 is important for GLR3.3 function. Moreover, all GLR3.3-VENUS fusion protein variants tested localized to the sieve element in punctate patterns as seen with the WT GLR3.3-VENUS fusion protein (Fig. 4*E*). Interestingly, GLR3.3 subcellular localization was also not affected by *isi1* mutation (Supplementary Fig. 6). These results are consistent, suggesting that the protein activity of GLR.3.3, rather than its subcellular localization, might be impaired by either *isi1* or RFL mutations. Given that ISI1 by itself does not have any apparent functional domain, it is considered less likely that ISI1 directly regulates GLR3.3 activity through RFL residues. We hypothesized that ISI1 possibly acts as a scaffold protein that links GLR3.3 to other regulators. Examples of scaffold proteins binding to members of the glutamate receptor superfamily exist. For instance, in plants, the animal homologous cornichon proteins CNIH1 and CNIH4 interact GLR3.3, controlling its membrane localization in pollen tubes (5). In animals filed, the C-tails of GluN2 NMDARs subunits end in a small motif (xSxV) that binds scaffold proteins regulating receptor localization in postsynaptic membranes (27, 34). It will be interesting to test this scaffold hypothesis. However, the regulation of GLR3.3 through RFL residues binding to ISI1 clearly represents a different kind of mechanism, as the subcellular localization of GLR3.3 is not affected by either RFL mutation in *GLR3.3* or *ISI1* loss-of-function.

In summary, we show in this work that the short C-tail of GLR3.3 is a key element in regulating its function in propagating long distance signals. We further identified ISI1 as a new interactor for GLR3.3. ISI1 was also found to be involved in multiple wound-induced responses. Moreover, three residues RFL in GLR3.3 C-tail were identified to be essential for binding to ISI1 and GLR3.3 function. Since a (modified) SWP still occurs in the absence of ISI1, these three residues may have further essential roles. While considerable research has focused on the channel properties of GLR3.3, our results isolate key functional C-tail residues and highlight from molecular and cellular perspectives the importance of the small C-tail of GLR3.3 in its actions.

## Materials and Methods

Materials and Methods are available in the Supplementary Information.

## Data Availability

RNAseq data have been deposited in the Gene Expression Omibus (GEO) (https://www.ncbi.nlm.nih.gov/geo/) under accession number GSE157938. All study data are included in the article, supplementary information and supplementary dataset 1.

## Acknowledgments

We would like to thank Prof. J. A Feijó for sharing the GLR3.3 cDNA, Dr. C.T. Nyguen for help with confocal microscopy, Lausanne Genomic Technologies Facility for RNA-seq service, and the Lausanne Cellular Imaging Facility. This work was supported by Swiss National Science Foundation grants 31003A-155960 and 3100A3-175566/1 to E.E.F.

## References

1. S. De Bortoli, E. Teardo, I. Szabo, T. Morosinotto, A. Alboresi, Evolutionary insight into the ionotropic glutamate receptor superfamily of photosynthetic organisms. Biophys Chem 218, 14–26 (2016).

2. M. M. Wudick, E. Michard, C. Oliveira Nunes, J. A. Feijo, Comparing Plant and Animal Glutamate Receptors: Common Traits but Different Fates? J Exp Bot 69, 4151–4163 (2018).

3. C. Ortiz-Ramirez et al., GLUTAMATE RECEPTOR-LIKE channels are essential for chemotaxis and reproduction in mosses. Nature 549, 91–95 (2017).

4. E. Michard et al., Glutamate receptor-like genes form Ca^2+^ channels in pollen tubes and are regulated by pistil D-serine. Science 332, 434–437 (2011).

5. M. M. Wudick et al., CORNICHON sorting and regulation of GLR channels underlie pollen tube Ca^2+^ homeostasis. Science 360, 533–536 (2018).

6. E. D. Vincill, A. E. Clarin, J. N. Molenda, E. P. Spalding, Interacting glutamate receptor-like proteins in Phloem regulate lateral root initiation in *Arabidopsis*. Plant Cell 25, 1304–1313 (2013).

7. H. Manzoor et al., Involvement of the glutamate receptor AtGLR3.3 in plant defense signaling and resistance to *Hyaloperonospora arabidopsidis*. Plant J 76, 466–480 (2013).

8. S. A. Mousavi, A. Chauvin, F. Pascaud, S. Kellenberger, E. E. Farmer, *GLUTAMATE RECEPTOR-LIKE* genes mediate leaf-to-leaf wound signalling. Nature 500, 422–426 (2013).

9. J. Browse, Jasmonate passes muster: a receptor and targets for the defense hormone. Annu Rev Plant Biol 60, 183–205 (2009).

10. M. Erb, P. Reymond, Molecular Interactions Between Plants and Insect Herbivores. Annual Review of Plant Biology 70, 527–557 (2019).

11. W. Mou et al., Ethylene-independent signaling by the ethylene precursor ACC in *Arabidopsis* ovular pollen tube attraction. Nat Commun 11, 4082 (2020).

12. Q. L. Shao, Q. F. Gao, D. Lhamo, H. S. Zhang, S. Luan, Two glutamate- and pH-regulated Ca^2+^ channels are required for systemic wound signaling in *Arabidopsis*. Sci Signal 13, eaba1453 (2020).

13. Z. Qi, N. R. Stephens, E. P. Spalding, Calcium entry mediated by GLR3.3, an Arabidopsis glutamate receptor with a broad agonist profile. Plant Physiol 142, 963–971 (2006).

14. A. Alfieri et al., The structural bases for agonist diversity in an *Arabidopsis thaliana* glutamate receptor-like channel. Proc Natl Acad Sci U S A 117, 752–760 (2020).

15. M. Weiland, S. Mancuso, F. Baluska, Signalling via glutamate and GLRs in *Arabidopsis thaliana*. Funct Plant Biol 43, 1–25 (2015).

16. P. H. Wang et al., The Glutamate Receptor-Like Protein GLR3.7 Interacts With 14-3-3omega and Participates in Salt Stress Response in *Arabidopsis thaliana*. Front Plant Sci 10, 1169 (2019).

17. I. F. Chang et al., Proteomic profiling of tandem affinity purified 14-3-3 protein complexes in *Arabidopsis thaliana*. Proteomics 9, 2967–2985 (2009).

18. T. Nagai et al., A variant of yellow fluorescent protein with fast and efficient maturation for cell-biological applications. Nat Biotechnol 20, 87–90 (2002).

19. C. T. Nguyen, A. Kurenda, S. Stolz, A. Chetelat, E. E. Farmer, Identification of cell populations necessary for leaf-to-leaf electrical signaling in a wounded plant. P Natl Acad Sci USA 115, 10178–10183 (2018).

20. A. Kurenda, E. E. Farmer, Rapid extraction of living primary veins from the leaves of *Arabidopsis thaliana*. Protocol Exchange doi:10.1038/protex.2018.119 (2018).

21. F. Rook et al., *Impaired sucrose induction1* encodes a conserved plant-specific protein that couples carbohydrate availability to gene expression and plant growth. Plant J 46, 1045–1058 (2006).

22. V. Salvador-Recatala, W. F. Tjallingii, E. E. Farmer, Real-time, *in vivo* intracellular recordings of caterpillar-induced depolarization waves in sieve elements using aphid electrodes. New Phytol 203, 674–684 (2014).

23. A. Kumari, A. Chetelat, C. T. Nguyen, E. E. Farmer, Arabidopsis H^+^-ATPase AHA1 controls slow wave potential duration and wound-response jasmonate pathway activation. Proc Natl Acad Sci U S A 116, 20226–20231 (2019).

24. G. L. Fernandez-Milmanda et al., A light-dependent molecular link between competition cues and defence responses in plants. Nat Plants 6, 223–230 (2020).

25. H. Husi, S. G. Grant, Isolation of 2000-kDa complexes of N-methyl-D-aspartate receptor and postsynaptic density 95 from mouse brain. J Neurochem 77, 281–291 (2001).

26. M. O. Collins, S. G. Grant, Supramolecular signalling complexes in the nervous system. Subcell Biochem 43, 185–207 (2007).

27. C. G. Lau, R. S. Zukin, NMDA receptor trafficking in synaptic plasticity and neuropsychiatric disorders. Nat Rev Neurosci 8, 413–426 (2007).

28. G. Hardingham, NMDA receptor C-terminal domain signaling in health and disease. Journal of Neurochemistry 150, 26–26 (2019).

29. L. Zhou, J. J. Du, The C-terminus of NMDAR GluN1-1a Subunit Translocates to Nucleus and Regulates Synaptic Function. Front Cell Neurosci 12 (2018).

30. X. Q. Hong et al., A novel function for the ER retention signals in the C-terminus of kainate receptor subunit, GluK5. Bba-Mol Cell Res 1866, 459–473 (2019).

31. R. Shin, J. M. Jez, A. Basra, B. Zhang, D. P. Schachtman, 14-3-3 proteins fine-tune plant nutrient metabolism. FEBS Lett 585, 143–147 (2011).

32. R. Hedrich, V. Salvador-Recatala, I. Dreyer, Electrical Wiring and Long-Distance Plant Communication. Trends Plant Sci 21, 376–387 (2016).

33. F. Rook et al., Impaired sucrose-induction mutants reveal the modulation of sugar-induced starch biosynthetic gene expression by abscisic acid signalling. Plant J 26, 421433 (2001).

34. L. Bard, L. Groc, Glutamate receptor dynamics and protein interaction: lessons from the NMDA receptor. Mol Cell Neurosci 48, 298–307 (2011).

