## Supplementary information for "A triresidue motif in the GLUTAMATE RECEPTOR-LIKE 3.3 C-tail interacts with IMPAIRED SUCROSE INDUCTION 1 and controls long distance wound signaling"

**This PDF file includes:**

Materials and Methods

Supplementary Figures 1 to 6

Supplementary Table 1

SI References

**Other supplementary materials for this manuscript include the following:**

Supplementary Dataset 1

### Materials and Methods

**Plant Materials and Growth Conditions.** *Arabidopsis* Col-0 was used as wild type and is the background of all the mutants investigated in this study. The T-DNA insertion lines *glr3.3a* (SALK_099757) and *glr3.6a* (SALK_091801) were reported in (1). *isi1-2* (SALK_014032) and *isi1-3* (SALK_045849) were from the Nottingham *Arabidopsis* Stock Center (NASC). Primers used for genotyping *ISI1* alleles are listed in the Supplementary Table 1. To work with 5-week-old plants, seeds were sown individually in 7 cm diameter pots. Plants were stratified at 4 °C for 2 days in the dark before transferring them to the growth room at 21°C under 150 $\mu E\ m^{-2}\ s^{-1}$ light (10 h light, 14 h dark, 70% humidity).

To generate *ISI1*$_{pro}$*:ISI1-mCherry/isi1-2* complementary lines, the full-length *ISI1* genomic sequence spanning 1175 bp promoter region and the gene (*ISI1*$_{pro}$*:ISI1*) was amplified and subsequently cloned into pUC57-L4-*Kpn1*/*Xma1*-R1 by digestion and ligation. Primers used for cloning *ISI1* are listed in the Supplementary Table 1. To obtain an *ISI1*$_{pro}$*:ISI1-mCherry* expression clone, *ISI1*$_{pro}$*:ISI1* in pUC57, *mCherry* coding sequence in L1-pDONOR221-L2 and the destination vector pEDO097pFR7m24GW (2) were combined by double Gateway cloning. Transgenic plants were obtained by dipping *isi1-2* plants with *Agrobacterium* carrying the corresponding vector. Similarly, translational reporter plants *IS11*$_{pro}$*:ISI1-GUSPlus* were made. In the latter case, mCherry in L1-pDONOR221-L2 was replaced by pEN-L1-GUS*Plus*-L2 for the final recombination, and Col-0 plants were transformed. The resulting T1 plants were selected based on seed coat fluorescence using a MZ16 FA microscope (Leica, Wetzlar, Germany). Homozygous T3 plants were used for all the analysis in this work.

To make *GLR3.3*$_{pro}$*:GLR3.3ΔCT-Venus/3.3a* plants, the *GLR3.3*$_{pro}$*:GLR3.3ΔCT* genomic fragment was amplified from the plasmid pUC57-*GLR3.3*$_{pro}$*:GLR3.3* genomic clone published in (3) and cloned into pUC57-L4-*Kpn1*/*Xma1*-R1 via Kpn1 and XmaI1 sites. Double Gateway cloning was carried out to combine *GLR3.3*$_{pro}$*:GLR3.3ΔCT* in pUC57 and pEN-L1-VENUS-L2 with the destination vector pH7m24GW. To introduce desired point mutations into the GLR3.3$_{pro}$:GLR3.3-VENUS fusions, PCRs were performed with mutagenic overlapping primers designed with the QuikChange Primer Design tool (https://www.agilent.com/store/primerDesignProgram.jsp) to amplify the entire plasmid pUC57-*GLR*$_{pro}$*:GLR* genomic. Primers used to generate the mutations are listed in the Supplementary Table 1. Together with pEN-L1-VENUS-L2, all the resulting pUC57-GLR3.3$_{pro}$:GLR3.3 clones with the indicated mutations were recombined with the destination vector pH7m24GW respectively to generate binary expression vectors and then transformed into *glr3.3a* mutant plants. T1 seeds were selected by adding 25 mg/mL hygromycin to the half-strength MS plates. T3 plants that were homozygous for the antibiotic were used for studies. At least two independent transgenic lines were used for the experiments in this work.

**Surface Potential Measurements.** Protocols for monitoring surface potential changes were detailed in (1, 3). Briefly, silver/silver chloride electrodes were placed on the petioles of both leaf 8 and 13 from 5-week-old plants. The connection between electrodes and leaf surface was maintained by adding one drop of 10 mM KCl in 0.5% (w/v) agar. A reference electrode was placed in the soil. For wounding, 50%-60% of the apical lamina surface distal to the rosette center of leaf 8 was crushed with a plastic forceps. Electrical signals were recorded from both leaves at 100Hz using LabScribe3 (iWorx System, Inc., Dover, NH) software. Amplitudes and durations of the measured electrical signals were analyzed as described in (1).

**Visualization of Protein Subcellular Localization by Confocal Microscopy**. To observe the subcellular localization of GLR3.3-VENUS and its derivatives, vein samples were prepared from expanded leaves of 5-week-old plants. Vein extraction was performed following description in (4). Isolated veins were immediately fixed with 4% paraformaldehyde (PFA) solution for 1 hour with gentle shaking, and then subjected to ClearSee treatment (5) for two days. Refreshing the ClearSee solution is necessary to get sufficiently cleared samples. To stain samples, 0.1% (w/v) Calcofluor-white was added into the ClearSee solution, and the samples were stained for 15 min. Then the samples were washed twice with ClearSee solution before observation. All the samples were visualized with a SP8 microscope (Leica Microsystems CMS GmbH, Mannheim, Germany). Sequential scanning mode was used to avoid interference between channels. VENUS was excited at 514 nm and detected in a range of 520-540 nm. In most cases, chlorophyll autofluorescence still remained and was detected in an emission window of 650-700 nm. Calcofluor-white was imaged with 405 nm excitation and 430-460 nm emission.

**Yeast Two-Hybrid Assay**. To fish potential interactors of GLR3.3, a yeast two-hybrid based screen was carried out with the ULTImate Y2H platform (Hybrigenics Services, Evry, France). For this, the C-terminal tail of GLR3.3 (850-933 aa) was constructed into pB27 vector (N-LexA-bait-C fusion) as a bait to screen against the prey cDNA library made from *Arabidopsis* rosette leaves. 146 millions interactions were analyzed. 84 clones were further processed. ISI1 appeared as one of the clones with very high confidence in the interactions. To further confirm the interaction between the C-tail of GLR3.3 and ISI1, the C-tail of GLR3.3 was cloned into the commercial pGBKT7 vector as bait, and ISI1 was inserted into pGADT7-Rec vector as prey. As controls, the C-tail of GLR3.1 and GLR3.6 were also constructed into pGBKT7 vector and tested their interactions with ISI1. Then the bait and prey constructs were co-transformed into the yeast strain AH109 and the interactions were analyzed following the protocol described in (6). To map the essential interacting sites in the C-tail of GLR3.3 as well as in ISI1, serial truncations and point mutation proteins were generated from the pGBKT7-3.3CT and pGAD-ISI1 plasmid templates respectively as described above using mutagenic overlapping primers. The primers used above are available in the *SI Appendix*, Table S1. The resulting bait and prey pairs were co-transformed into yeast cells and the interactions were analyzed by growing transformants on selective medium.

**Firefly Luciferase Complementation Imaging (LCI) Assay.** The full-length *GLR3.3* cDNA (7) was fused upstream of the N-terminal part of *Luciferase* (*nLUC*) in the pCAMBIA1300-nLUC vector by infusion cloning. Similarly the C-tail of *GLR3.3* was introduced by conventional cloning via sites Kpn1 and Sal1. *ISI1* cDNA was fused downstream of the C-terminal part of *Luciferase* (*cLUC*) in the pCAMBIA1300-cLUC vector. The primers used for the above constructions are listed in the *SI Appendix*, Table S1. All the resulting constructs along with the empty vectors were transferred into *Agrobacterium* strain GV3101, respectively. To determine the interactions of full length or the C-tail GLR3.3 with ISI1 in *Nicotiana benthamiana* leaves, *Agrobacterium* harboring the indicated constructs were resuspended in the infiltration buffer containing 10 mM $MgCl_2$, 10 mM MES, 0.5 g/L glucose and 150 μM acetosyringone to a final concentration of $OD_{600}$=0.5. Then equal volumes of different combinations were mixed and coinfiltrated into the abaxial face of tobacco leaves using a needless syringe. Plants were then kept for 24 h in the dark before transferring them to light for another 24 to 48 h. To facilitate and observe the luminescence brought about by the interactions of the proteins, the *N. benthamiana* leaves were fully infiltrated with 0.1 mg/mL luciferin and placed in the dark for 5 min before CCD imaging. LUC activity was determined by IVIS Lumina III In Vivo Imaging System (PerkinElmer, Richmond, CA). The exposure time was from 1 to 5 mins depending on the signal intensity.

**GUS Staining and Sectioning**. Three-week-old *IS11$_{pro}$:ISI1-GUSPlus/Col0* plants were excised and immediately fixed with 90% acetone on ice for 1 hour, followed by twice washing with 50 mM sodium phosphate buffer (pH 7.4). Then the plants were stained by adding staining solution (10 mM $Na_2EDTA$, 50 mM sodium phosphate buffer, 1 mM $K_4Fe(CN)_6$, 1 mM $K_3Fe(CN)_6$, 0.1% (v/v) Triton X-100, 0.5 mg/mL X-Gluc (pH 7.2) ) and subjected to vacuum infiltration for 30 min. After incubating at 37 °C in the dark for 6 hours, the plants were washed with 50 mM sodium phosphate buffer and then cleared with 70% (v/v) ethanol. Images of plants were taken with a VHX-6000 digital microscope (Keyence, Osaka, Japan). To study the detailed expression pattern of ISI1 at cellular level, the petioles of the expanded leaves were further fixed in glutaradehyde/formaldehyde/50 mM sodium phosphate (pH 7.2) 2:5:43 (v/v/v) for 30 min, and then dehydrated with ethanol gradients (10%, 30%, 50%, 70% 90% and twice absolute) for 30 min in each concentration. Afterwards they were embedded in Technovit 7100 resin (Haslab GmbH, Ostermundigen, Switzerland) according to the manufacturer's instructions. Transversal sections (5 μM thick) were made on a RM2255 microtome (Leica, Wetzlar, Germany). The sections were mounted in 40% (v/v) glycerol and then imaged with a Leica DM5500 microscope.

**Protein Extraction and Western Blot**. Approximately 100 mg midveins from 5-6 week-old *$ISI1_{pro}$:ISI1-mCherry/isi1-2* plants were harvested according to protocol in (4) and frozen for protein extraction. The lamina parts of the leaves after midvein removal were also collected for analysis. The frozen tissues were ground to fine powder with a TissueLyser (Qiagen, Hilden, Germany). Proteins were extracted with lysis buffer (50 mM Tris-HCl, pH 7.5, 150 mM NaCl, 0.1% (v/v) Nonidet P-40) plus plant-specific protease inhibitor cocktail (Sigma-Aldrich Chemie GmbH, Buchs SG, Switzerland). After centrifuging at 13,000 rpm for 15 min at 4 °C, the supernatants were collected. The protein samples were prepared by mixing the supernatant with 4 × SDS protein loading buffer, followed by incubating for 5 min at 95 °C, and then separated by running a 4%-12% (v/v) polyacrylamide gradient ExpressPlus™ Bis-Tris gel (GenScript, Piscataway, NJ). Immunoblotting was employed to detect the ISI1-mCherry fusion protein by using anti-mCherry antibody (ab167453, polyclonal, Abcam, Cambridge, UK). Ponceau staining was used to assess correct gel loading.

**RNA Extraction and RT-qPCR.** Wounding experiments for *JAZ10* RT-qPCR were performed with 5-week-old plants. The procedures for cDNA reverse transcription and quantitative PCR were described in (8). q-PCR data were normalized to the reference gene ubiquitin-conjugating enzyme 21 (UBC21). Primers for *UBC21* and *JAZ10* were used previously (1). Primers to detect transcripts generated from the 5' end and 3'end of *ISI1* as shown in Supplementary Fig. 1A are listed in the Supplementary Table 1.

**RNA Sequencing.** 5-week-old Col-0 and *isi1-2* plants were used. Distal leaves 13 were harvested one hour after wounding leaves 8. As controls, leaves 13 from unwounded plants were also collected. Two individual plants were pooled for one replicate and three replicates were used. Total RNAs from all samples were purified with ReliaPrep™ RNA Tissue Miniprep System (Promega, Madison, WI). RNA qualities were assessed with a Fragment Analyzer (Agilent, Santa Clara, CA). Truseq Stranded mRNA kits (Illumina, San Diego, CA) were used to prepare cDNA libraries, which were then sequenced (single end) on an Illumina Hiseq 4000 instrument. Reads were aligned against *Arabidopsis thaliana* TAIR10.39 genome using STAR (v.2.5.3a) (9). Differential expression was computed with limma (10). GO analysis (Gene classification and enrichment analysis) were performed with the clusterProfiler package (11) (v. 3.12.0) in R (v. 3.3.1) for the GO term biological process.

**Electrical Penetration Graph (EPG) Recordings.** EPG was employed to study sieve elements-specific electrical signals. The experimental setup and data analysis were detailed previously (12, 13).

**Jasmonate Quantifications.** 5-week-old plants were used. Jasmonates were measured in wounded and distal leaves at different time points. At least three biological replicates were analyzed. Hormone extraction and quantification were following the protocol in (14).

**Insect Bioassays.** Eleven pots of 5-week-old plants were placed in Plexiglass boxes (28.5×19×19 cm). Four freshly hatched *Spodoptera littoralis* larvae were gently placed on the rosette center of each plant with a soft brush. After feeding for up to 10 days, the caterpillars were collected and weighed from individual boxes. The caterpillar mass from each box was considered as one replicate. The average weight from four replicates and the total numbers of the surviving caterpillars were recorded.

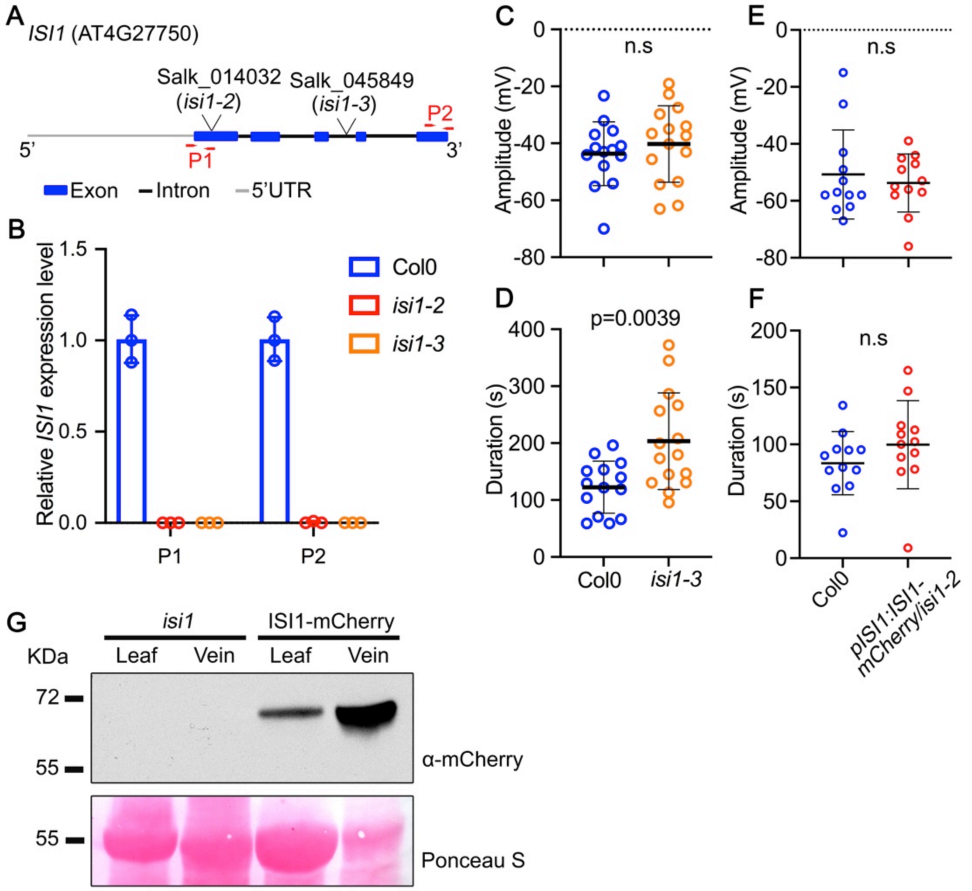


**Supplementary Fig. 1. Wound-related phenotypes of a second allele of *isi1 and isi1-2* complementary line.** (**A**) The *ISI1* gene model. The positions of the T-DNAs from two *isi1* alleles are shown. (**B**) *ISI1* transcript levels in unwounded leaves from wild type and *isi* mutants. P1 and P2 in (**A** and **B**) indicate the regions that are amplified. Circles represent individual values. Data shown are means ± SD. n=3. (**C** and **D**) Quantification of the amplitudes (**C**) and durations (**D**) of surface potential changes in the distal leaves of *isi1-3* compared to wild type plants. n.s, not significant. Colored circles shown are measurements from individual plants. n=14-15. *p* values were calculated with two tailed Student's *t*-tests. (**E** and **F**) Amplitudes and durations recorded in leaf 13 from Col-0 and $ISI1_{pro}$*:ISI1-mCherry/isi1-2* plants after wounding leaf 8. Circles represent individual biological replicates. Data shown are means ± SD. n=12. (**G**) Western blot for ISI-mCherry fusion proteins from midveins or lamina tissues from which the midveins were removed using protocol in (4). Proteins were extracted from 5-week-old $ISI1_{pro}$*:ISI1-mCherry* plants and detected with mCherry antibodies. Ponceau S staining was used to assure comparable loading of each well.

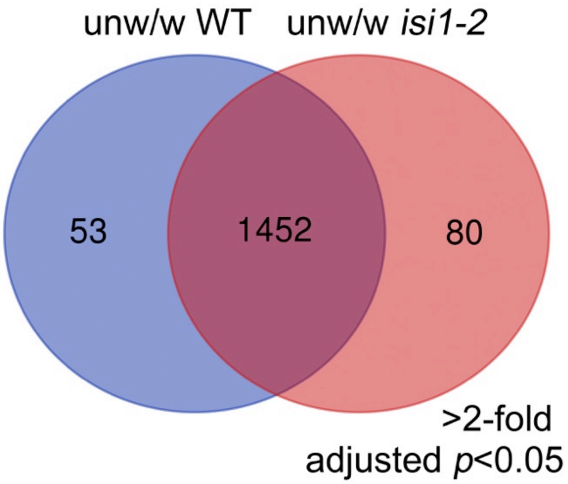


**Supplementary Fig. 2. Comparison of differentially expressed wound-induced genes in WT and *isi1-2* from RNA-seq.** Samples were harvested from distal leaf 13 one hour after wounding leaf 8 of both wild type and *isi1-2* mutant plants. The venn diagram depicts the numbers of genes that were upregulated more than twofold after wounding. Adjusted *p* values<0.05.

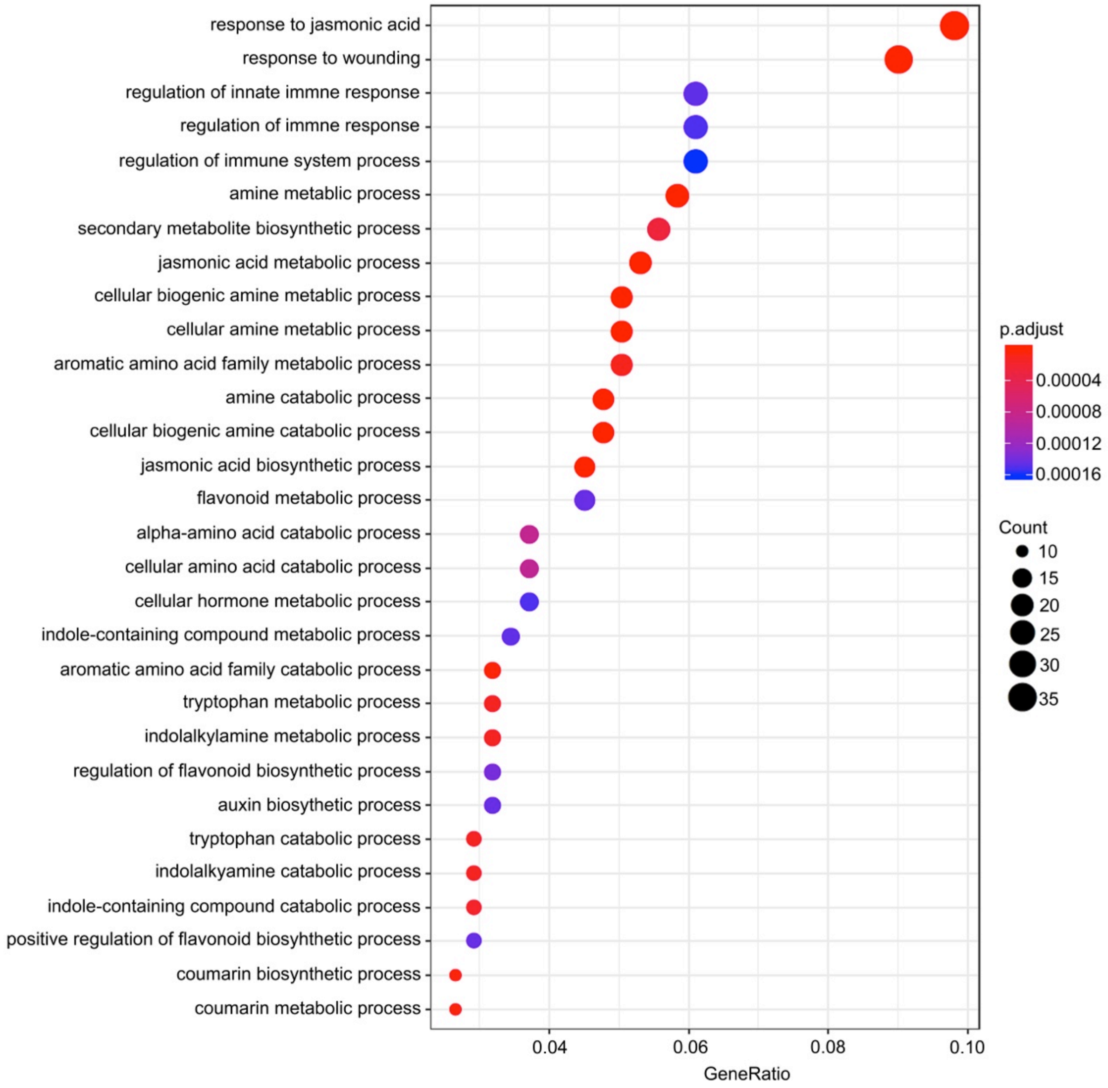


**Supplementary Fig. 3. Gene Ontology enrichment analysis showing genes upregulated in wounded *isi1-2* plants in comparison to WT.** The enrichment was analyzed for the GO term biological processes. Only results giving significantly enriched gene sets are presented. GeneRatio is the percentage of total differentially expressed genes (DEGs) for the given GO term. Statistics were calculated with Benjamini-Hochberg method; adjusted $p$ values $<0.05$.

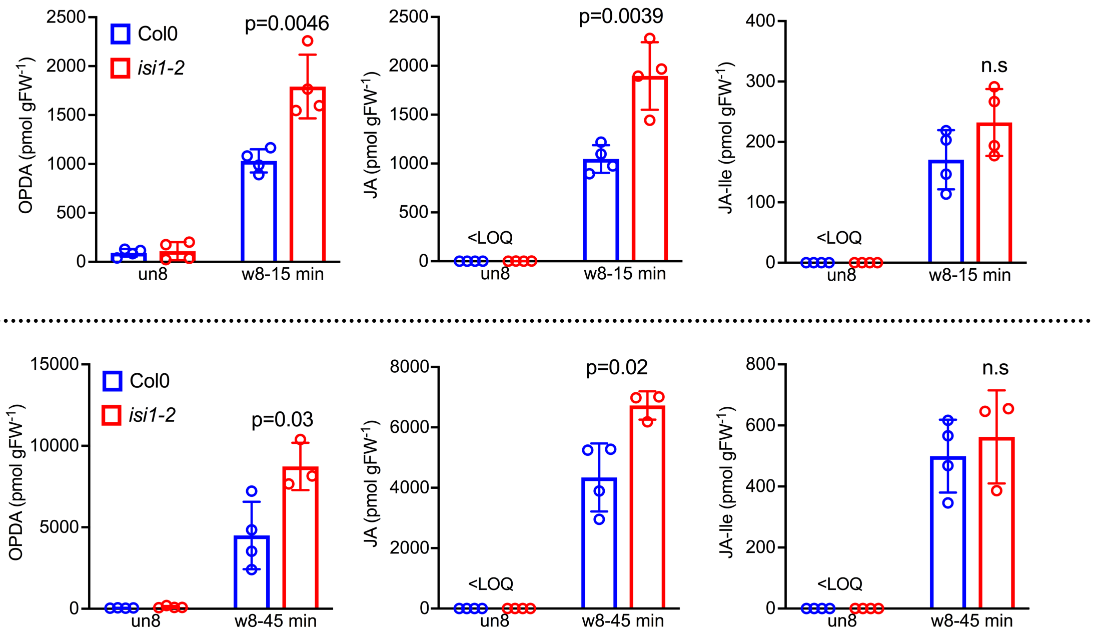


**Supplementary Fig. 4. Jasmonate accumulation in the *isi1-2* mutant after wounding.** Levels of OPDA, JA and JA-Ile were measured in both unwounded leaf 8 (un8) and wounded leaf 8 (w8) at different time points (15 min and 45 min) post wounding. In each case at least three biological replicates were analyzed. Data shown are means ± SD. *p* values were calculated by two tailed Student's *t*-tests. n.s, not significant. LOQ: limit of quantification.

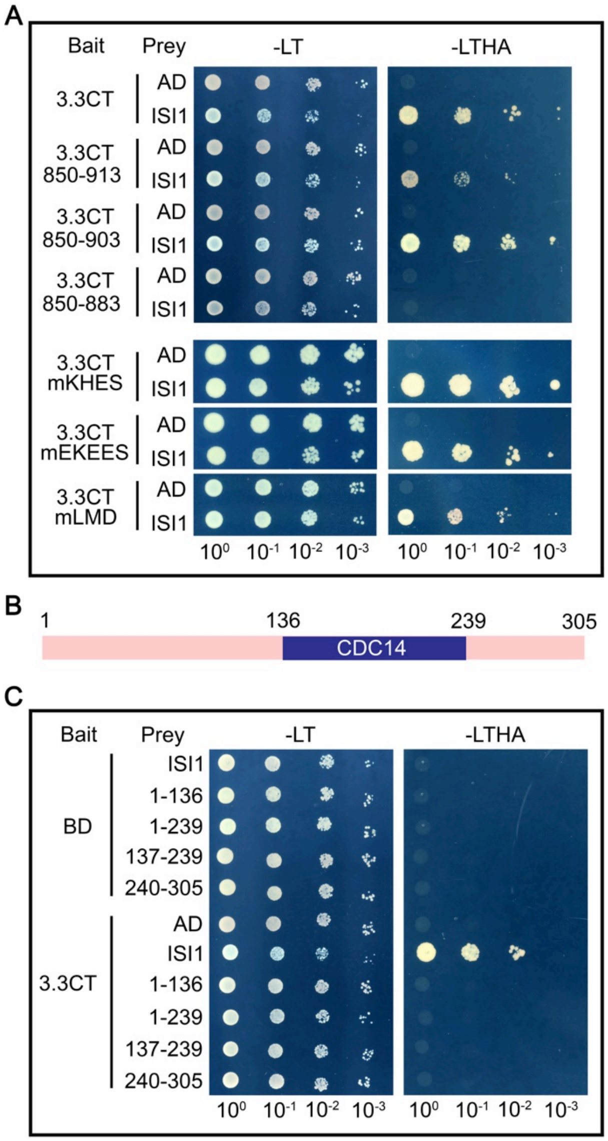


**Supplementary Fig. 5. Domain mapping for GLR3.3-ISI1 binding sites in Y2H assays.** (**A**) ISI1 interaction with GLR3.3 C-tail (CT) variants. Images were taken after 3 or 5 days for yeast groups that were grown on Leu-Trp- (-LT)/Yeast Nitrogen Base (YNB) or Leu-Trp-Histidine-Adenine (-LTHA)/YNB medium, respectively. GLR3.3 CT and its mutants were co-expressed with empty AD (Activating Domain) vectors as negative controls. All the GLR3.3 C-tail variants (deletions and point mutations) were generated by PCRs using overlapping mutagenic primers and Binding Domain (BD)-3.3CT plasmid as template. (**B**) Schematic model for ISI1 protein. Residues from 136 to 239 in ISI1 were predicted by Pfam database to have a homology to Cell Division Control protein 14 (CDC14) in fission yeast. (**C**) Interaction analysis for the entire GLR3.3 C-tail with different ISI1 truncations. Empty BD vectors co-transformed with ISI1 truncated proteins served as negative controls.

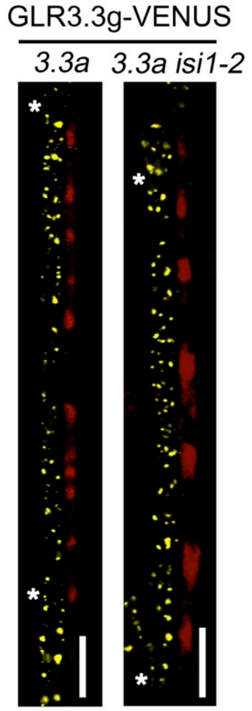


**Supplementary Fig. 6. Subcellular localization of GLR3.3 is not affected by ISI1 inactivation.** GLR3.3-VENUS localization in the *isi1-2* background. To study the impact of *ISI1* mutation on GLR3.3 intracellular distribution, the *glr3.3a* mutant complemented with full-length *GLR3.3* genomic sequence (*GLR3.3g*) from (3) was crossed with *isi1-2*. VENUS fluorescence was detected by confocal microscopy. Asterisks indicate the positions of sieve plates. Bars in all the images =10 μm. Images were taken under the same parameters in each panel.

**Supplementary Table 1.** Primer list in this study.

**Primers used for genotyping T-DNA mutants**

| Primer name | Primer sequence (5’-3’) |
|---|---|
| ISI1-2-RP<br>ISI1-2-LP | CATCAGCCATTTTTCCAGTTC<br>CACCTCTCACTCTGATTTGGC |
| ISI1-3-RP<br>ISI1-3-LP | CTCCATCCTCAGAGCACTGTC<br>GGACACGTAAATGGGAAGGAT |

**Primers used for cloning**

| Primer name | Primer sequence (5’-3’) | Vectors |
|---|---|---|
| ISI1(Pro+Gene) -F<br>ISI1(Pro+Gene) –R | ggaattcGGTACCATGTATCTGAAGAGACCGATATGG<br>ttcccccccgggagtacaagtcaagagactcgagca | pUC57 |
| GLR3.3-S-Kpn1infusion<br>GLR3.3-A-Sal1infusion | CGGGGGACGAGCTCGGTACCATGAAGCAACTCTGGACTTT<br>ACGAGATCTGGTCGACGTCTAATGGATTTACCGAATT | pCAMBIA1300-nLUC |
| 3.3CT-S-Kpn1<br>3.3CT-2A-Sal1 | cggGGTACCATGCAGATCATCCGTCAGCTCT<br>gcGTCGACGTCTAATGGATTTACCGAATT | pCAMBIA1300-nLUC |
| ISI1-S-kpn 1<br>ISI1-A-Sal1 | ggaattcGGTACCATGTATCTGAAGAGACCGATATGG<br>acgcGTCGACTTAGTACAAGTCAAGAGACTCGAGC | pCAMBIA1300-cLUC |
| 3.3CT-S-EcoR1<br>3.3CT-A-Sal 1 | cgGAATTCCAGATCATCCGTCAGCTCTATA<br>gcGTCGACTCAGTCTAATGGATTTACCGA | pGBKT7 |
| 3.6CT-S-EcoR1<br>3.6CT-A-Sal 1 | cgGAATTCCGTCAGTTTGGACAGCAATG<br>gcGTCGACTTAGTTGCAGCGACTTGAACC | pGBKT7 |
| 3.1CT-S-EcoR1<br>3.1CT-A-Sal 1 | cgGAATTCGTGCATAGCTTCTGGGGTATG<br>gcGTCGACTCATATGGGTCTTCTAGATGCAG | pGBKT7 |
| ISI1-S-Nde 1<br>ISI1-A-Sal 1 | ggaattcCATATGATGTATCTGAAGAGACCGATATGG<br>acgcGTCGACTTAGTACAAGTCAAGAGACTCGAGC | pGADT7-Rec |
| GLR3.3$_{pro}$:GLR3.3 ΔCT-F<br>GLR3.3$_{pro}$:GLR3.3 ΔCT-R | gtatagaaaagttg ggtacc acccaaaccgcttattcttg<br>tgtacaaacttgt cccggg aacaaagtataggaagag | pUC57 |

**Primers used for site-directed mutagenesis**

| Primer name | Primer sequence (5’-3’) |
|---|---|
| 3.3CT-mKKRK-F<br>3.3CT-mKKRK-R | tcccgatgtatcgttcattgaaccatcgatcgctgccgccgcgctttcgtgcttggactcttctttatcatcc<br>ggatgagaaagaagagtccaagcacgaaagcgcggcggcagcgatcgatggttcaatgaacgatacatcgg |

| | |
|---|---|
| | ga |
| 3.3CT-mS-F | ctcgtttgcaaagattcttggctctcatggatgagaaagaa |
| 3.3CT-mS-R | ttctttctcatccatgagagccaagaatctttgcaaacgag |
| 3.3CT-mRFL-F | ctcttctttctcatccatgagagacgcggctgcttgcaaacgagtggagcgcatggag |
| 3.3CT-mRFL-R | cctccatgcgctccactcgtttgcaagcagccgcgtctctcatggatgagaaagaaga |
| 3.3CT850-913-S | tgaacgatacatcgggatgagtcgacctgcag |
| 3.3CT850-913-A | ctgcaggtcgactcatcccgatgtatcgttca |
| 3.3CT850-903-S | cgctgcaggtcgactcactttctcttcttgcttt |
| 3.3CT850-903-A | aaagcaagaagagaaagtgagtcgacctgcagcg |
| 3.3CT850-883-S | caggtcgactcattgcaaacgagtggagcgca |
| 3.3CT850-883-A | tgcgctccactcgtttgcaatgagtcgacctg |
| 3.3CT-mLMD-S | gcttggactcttctttctcagccgcggcagacaagaatctttgcaaacgagtggagc |
| 3.3CT-mLMD-A | gctccactcgtttgcaaagattcttgtctgccgcggctgagaaagaagagtccaagc |
| 3.3CT-mEKEES-S | atctttctcttcttgctttcgtgcttggccgctgctgccgcatccatgagagacaagaatctttgcaa |
| 3.3CT-mEKEES-A | ttgcaaagattcttgtctctcatggatgcggcagcagcggccaagcacgaaagcaagaagagaaagat |
| 3.3CT-mKHES-S | gttcattgaaccatcgatctttctcttcttggctgcggccgcggactcttctttctcatccatgagagacaag |
| 3.3CT-mKHES-A | cttgtctctcatggatgagaaagaagagtccgcggccgcagccaagaagagaaagatcgatggttcaatgaac |
| 3.3CT-mR-S | atccatgagagacaagaatgcttgcaaacgagtggagcgc |
| 3.3CT-mR-A | gcgctccactcgtttgcaagcattcttgtctctcatggat |
| 3.3CT-mFL-S | cttctttctcatccatgagagacgcggctctttgcaaacgagtggagcgc |
| 3.3CT-mFL-A | tgcgctccactcgtttgcaaagagccgcgtctctcatggatgagaaagaag |
| ISI1-1-239-S | tgcggagagttcttactataagtcgagctgcagatg |
| ISI1-1-239-A | catctgcagctcgacttatagtaagaactctccgca |
| ISI1-1-136-S | tcagtgtggagccgatgaagatataagtcgagctgc |
| ISI1-1-136-A | gcagctcgacttatatcttcatcggctccacactga |

### Primers used for qPCR

| Primer name | Primer sequence (5'-3') |
|---|---|
| ISI1-RS1 | GAATCCATCAGAATCGGAGACC |
| ISI1-RA1 | CGGATGCGAAGGAAGGAGA |
| ISI1-RS2 | CAATAGCGAGTGTAAATGAAGACAT |
| ISI1-RA2 | TCTACCAGCCTGGATATGAAGC |
| ST2a-RS | GATCGAGAAACCCGGTGTGAA |
| ST2a -RA | CGATCACTCCCCGGCAATAC |

**Supplementary Dataset 1.** List for the potential GLR3.3 C-tail interactors from Y2H screen.

### SI References


1. S. A. Mousavi, A. Chauvin, F. Pascaud, S. Kellenberger, E. E. Farmer, *GLUTAMATE RECEPTOR-LIKE* genes mediate leaf-to-leaf wound signalling. *Nature* **500**, 422-426 (2013).
2. T. L. Shimada, T. Shimada, I. Hara-Nishimura, A rapid and non-destructive screenable marker, FAST, for identifying transformed seeds of *Arabidopsis thaliana*. *Plant J* **61**, 519-528 (2010).
3. C. T. Nguyen, A. Kurenda, S. Stolz, A. Chetelat, E. E. Farmer, Identification of cell populations necessary for leaf-to-leaf electrical signaling in a wounded plant. *P Natl Acad Sci USA* **115**, 10178-10183 (2018).
4. A. Kurenda, E. E. Farmer, Rapid extraction of living primary veins from the leaves of *Arabidopsis thaliana*. *Protocol Exchange* doi:10.1038/protex.2018.119 (2018).
5. R. Ursache, T. G. Andersen, P. Marhavy, N. Geldner, A protocol for combining fluorescent proteins with histological stains for diverse cell wall components. *Plant J* **93**, 399-412 (2018).
6. Q. Wu *et al.*, Ubiquitin Ligases RGLG1 and RGLG5 Regulate Abscisic Acid Signaling by Controlling the Turnover of Phosphatase PP2CA. *Plant Cell* **28**, 2178-2196 (2016).
7. M. M. Wudick *et al.*, CORNICHON sorting and regulation of GLR channels underlie pollen tube $Ca^{2+}$ homeostasis. *Science* **360**, 533-536 (2018).
8. A. Gfeller *et al.*, Jasmonate controls polypeptide patterning in undamaged tissue in wounded Arabidopsis leaves. *Plant Physiol* **156**, 1797-1807 (2011).
9. A. Dobin *et al.*, STAR: ultrafast universal RNA-seq aligner. *Bioinformatics* **29**, 15-21 (2013).
10. M. E. Ritchie *et al.*, limma powers differential expression analyses for RNA-sequencing and microarray studies. *Nucleic Acids Res* **43**, e47 (2015).
11. G. Yu, L. G. Wang, Y. Han, Q. Y. He, clusterProfiler: an R package for comparing biological themes among gene clusters. *OMICS* **16**, 284-287 (2012).
12. V. Salvador-Recatala, W. F. Tjallingii, E. E. Farmer, Real-time, *in vivo* intracellular recordings of caterpillar-induced depolarization waves in sieve elements using aphid electrodes. *New Phytol* **203**, 674-684 (2014).
13. A. Kumari, A. Chetelat, C. T. Nguyen, E. E. Farmer, *Arabidopsis* $H^+$-ATPase AHA1 controls slow wave potential duration and wound-response jasmonate pathway activation. *Proc Natl Acad Sci U S A* **116**, 20226-20231 (2019).

14. G. Glauser, A. Vallat, D. Balmer, Hormone profiling. *Methods Mol Biol* **1062**, 597-608 (2014).
